# Resilience of riparian vegetation composition and diversity following cessation of livestock grazing in northeastern Oregon

**DOI:** 10.1101/2021.04.08.438927

**Authors:** J Boone Kauffman, Greg Coleman, Nick Otting, Danna Lytjen, Dana Nagy, Robert L. Beschta

## Abstract

Riparian ecosystem restoration has been accomplished through exclusion of livestock using corridor fencing along hundreds of kilometers of streams in the western USA, for the benefit of riparian-obligate wildlife and endangered fishes. Yet few studies have evaluated shifts in the vegetation composition and diversity following the cessation of livestock impacts. We sampled riparian vegetation composition along 11-paired grazed and ungrazed (exclosed) stream reaches in northeastern Oregon, USA. Exclosure ages ranged from 2 to >30 years and grazing treatments varied from light grazing every one out of three years to heavy season-long grazing. Species richness and diversity was higher in the ungrazed reaches (p =0.002). The abundance of native sedges (*Carex* spp.) and broad-leaved forbs were also significantly (p < 0.05) greater in ungrazed areas. In contrast, exotic species adapted to grazing such as *Poa pratensis* and *Trifolium repens* were more abundant in grazed stream reaches. The prevalence of hydrophytic species significantly increased (p ≤ 0.01) in ungrazed reaches, (based on wetland species indicator scores), indicating that wetland-dominated communities within the ungrazed stream reaches were replacing ones adapted to drier environments. The increased abundance of facultative and wetland-obligate species in ungrazed reaches compared to grazed reaches suggests that livestock grazing exacerbates those climate change effects also leading to warmer and drier conditions. Further, riparian-obligate shrub cover along the streambank was higher in 7 of 8 exclosures that were older than 5 years. As a restoration approach the inherent resilience of riparian ecosystems exhibited in ungrazed riparian zones suggest positive feedbacks to other beneficial ecosystem processes such as increased species and habitat diversity, increased carbon sequestration, enhanced allochthonous inputs and greater sediment retention, that would affect the aquatic and terrestrial biota, water quality, and stream morphology.

## Introduction

Riparian areas are zones of contact between land and water ecosystems, represented by mesic, productive environments bordering streams, rivers, lakes, and springs [1]. Whereas riparian zones comprise only 1-2% of western USA landscapes, they provide habitat for more wildlife species than any other vegetation type [2] For example, about 70% of the wildlife species in the Pacific Northwest region, USA depend on riparian areas for all or part of their life cycle [3]. It has been estimated that 204 (77%) of the 266 species of inland birds that breed in Oregon and Washington do so in riparian and wetland environments [3].

Riparian vegetation is a keystone ecosystem feature that exerts a strong influence on adjacent uplands and aquatic ecosystems. Riparian plant communities are integral to stream function/aquatic productivity, especially in low-order streams where they strongly influence stream temperatures [4] channel form [5], and the habitats of fish and aquatic invertebrates [6,7] In addition, they are the predominant in-stream sources of nutrients and carbon via allochthonous inputs [7,8]. Through interactions of soil, vegetation, and water, riparian areas retain and filter sediments, stabilize stream banks, and moderate stream and groundwater flows through storage and flood attenuation. Productivity of riverine fish communities is determined by both habitat and food resources, factors that are intricately linked to the structure and composition of riparian zones [7].

Riparian zones and other palustrine/riverine wetlands are also important sinks of atmospheric carbon, underscoring their values for inclusion in climate change mitigation and adaptation strategies. Nahlik and Fennessy [9] reported that soils of palustrine/riverine wetlands of the USA had mean carbon stocks of 3687 Mg C ha^-1^ and that western wetlands stored 236 Mg C ha^-1^. These stocks are about 3 to 6 times that of adjacent upland forests (≈61 Mg C ha^-1^) of the Blue Mountains of Oregon (the location of this study; [10]).

Given the important ecosystem functions and ecological services provided by riparian vegetation, shifts in structure and composition would like have far reaching effects on both adjacent terrestrial and aquatic ecosystems. In the western USA, livestock grazing is the most widespread land use on public lands and has been suggested to be a significant influence affecting riparian ecosystem structure, diversity, and function [11-13]. Cattle tend to prefer and congregate in riparian areas because of the abundant forage, proximity to water, relatively level terrain, and favorable microclimate, thus causing substantial damage to stream and riparian ecosystems [11,12,14].

Livestock grazing has been the most prevalent cause of ecological degradation of riparian/stream ecosystems in the Intermountain west [11,15,16]. Elmore and Kauffman [12], Beschta et al. [17] and Kauffman et al. [18] suggested that the cessation of livestock grazing in riparian zones of eastern Oregon was the single most ecologically and economically effective approach for restoring salmonid habitats.

Several studies have examined effects of livestock effects on riparian vegetation including effects on root mass [19], wetland species abundance [20], and vegetation structure [21,22] Most studies that have quantified the vegetation differences between grazed and ungrazed stream reaches in a wide diversity of stream types in the Pacific Northwest have focused on the shrub component [21-23]. However, few studies have examined how livestock affect riparian composition as manifested in species diversity, and richness.

Kauffman et al. [24] suggested the first logical step in riparian restoration is the implementation of “passive restoration” defined as the cessation of those activities that are causing ecosystem degradation or preventing recovery. Cessation of livestock grazing via exclusion fencing along salmonid bearing streams has been a common passive restoration approach to fish and wildlife habitat restoration in the Interior Colombia Basin of Oregon and Washington. Because the vegetation of riparian zones are adapted to frequent fluvial disturbances [7], many species possess adaptations facilitating a rapid recovery following both natural and anthropogenic disturbances.

The objective of this study was to quantify changes in the composition and structure, of the riparian vegetation composition along 11 experimental streams where passive restoration (corridor fencing or livestock exclosures) had occurred. From measurements of species cover, we calculated composition, richness, and diversity in paired reaches that included livestock exclusion with an adjacent reach where the riparian zones were grazed by domestic cattle. We hypothesized that given the inherent resilience of riparian vegetation and physical shifts that occur due to livestock removal [19], the composition of ungrazed stream reaches would have a greater dominance of hydrophytic vegetation, as well as increased, species richness, and diversity.

## Methods

To examine how streamside riparian vegetation differed between grazed and ungrazed stream reaches a total of 11 Northeast Oregon streams were selected. These streams were all tributaries of the Columbia River (Fig 1). Land tenure consisted both of public and private ownership. Each study stream consisted of two reaches, a grazed reach and an exclosed (ungrazed) reach (Table 1). Grazed reaches were those in which livestock grazing (principally cattle) was a dominant use in the riparian zone and surrounding uplands. Exclosed reaches were those where livestock grazing had been eliminated through the construction of riparian exclosures or corridor fences. Riparian livestock exclosures are essential research tools for the study of ecosystem processes, recovery, and to better inform livestock management [25] A major advantage of using small exclosures is that environmental site variability (precipitation, geology, flow regime, etc.) is practically the same for adjacent grazed and ungrazed areas, thus isolating the potential influence of livestock (or livestock removal [20,25].

**Table 1.** Site characteristics of the 11 stream reaches selected for study in northeast Oregon, USA.

| Site | Period of<br>exclusion<br>(years) | Site elevation<br>(m) | Mean annual ppt<br>(in) | Drainage<br>area<br>(km<sup>2</sup>) | Sinuosity<br>(m/m) | Valley gradient<br>(%) | Channel<br>gradient<br>(%) |
| --- | --- | --- | --- | --- | --- | --- | --- |
| Bear Creek (Silvies) | 2 | 1554 | 27 | 39.2 | 1.55 | 0.99 | 0.64 |
| Camas Creek | 5 | 1240 | 25 | 95.3 | 1.22 | 0.72 | 0.58 |
| Chesnimnus Creek | 14 | 1305 | 19 | 40.5 | 1.27 | 1.77 | 1.39 |
| Camp Creek | 36 | 1467 | 25 | 16.4 | 1.33 | 3.43 | 2.58 |
| Devil's Run Creek | 10 | 1285 | 19 | 29.0 | 1.75 | 1.86 | 1.05 |
| Middle Fk John Day | 3 | 1292 | 21 | 97.6 | 1.91 | 0.43 | 0.23 |
| Murderers Creek | 30 | 1347 | 19 | 36.3 | 1.58 | 0.52 | 0.34 |
| Summit Creek | 22 | 1506 | 23 | 77.3 | 1.45 | 0.90 | 0.62 |
| Lower Swamp Creek | 13 | 1123 | 19 | 79.9 | 1.29 | 0.60 | 0.46 |
| Upper Swamp Creek | 13 | 1142 | 19 | 74.6 | 1.33 | 0.77 | 0.56 |
| Tex Creek | 23 | 1359 | 19 | 31.6 | 1.24 | 1.11 | 0.90 |

**Figure 1.**
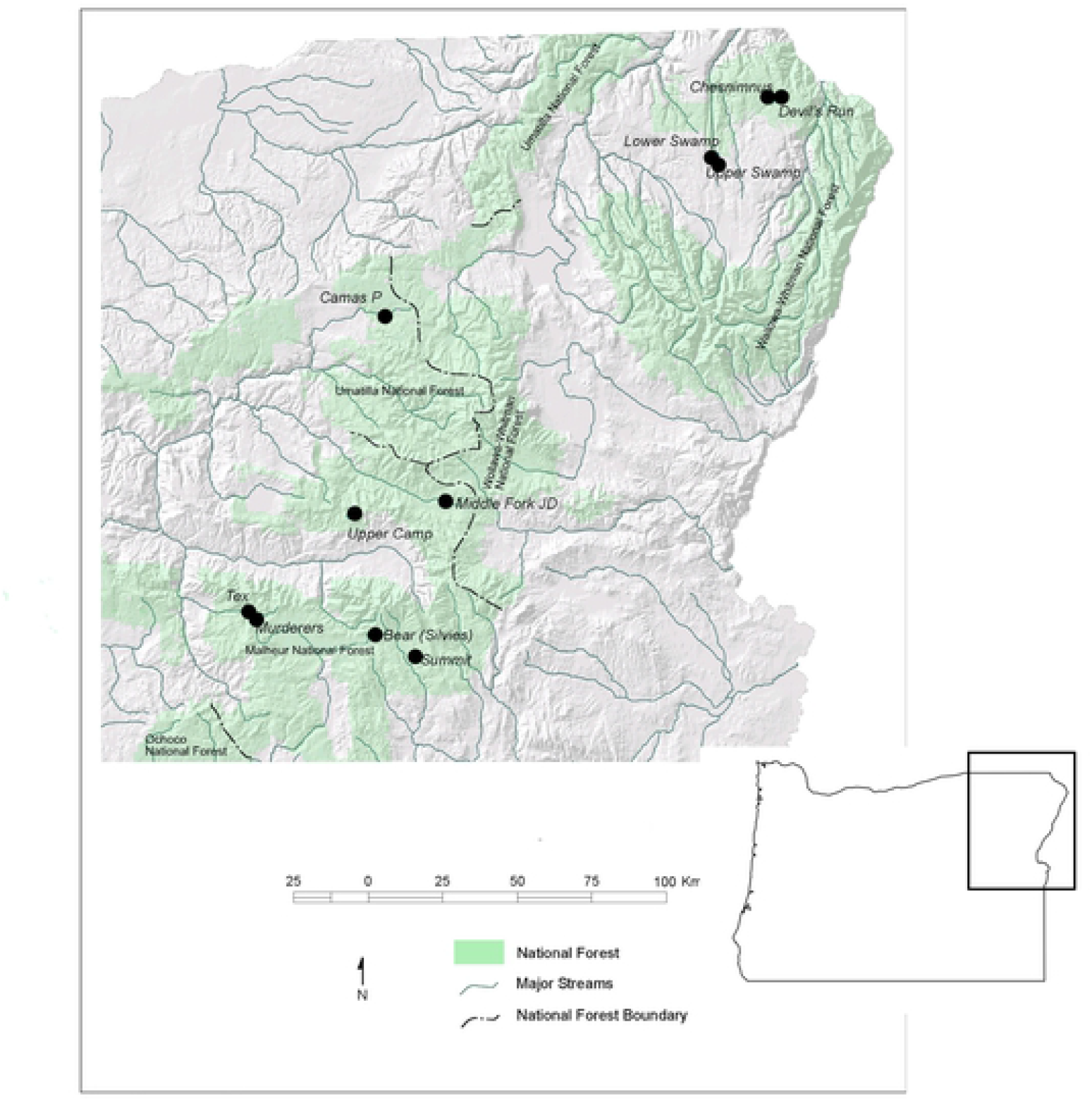
Location of the 11 study stream reaches selected for study in Northeastern Oregon, USA

Exclosure ages ranged from ∼2 to 37 years (Table 1). Criteria for site selection included paired reaches (grazed and ungrazed) that were as geomorphically similar as possible, streams with salmonids, knowledge of the history of the exclosure, and owner permission. Our assumptions were that prior to the construction of exclosure fences the vegetation composition was similar between reaches. The adjacent nature of sampled areas with similar geomorphic surfaces increased the likelihood that differences between grazed and ungrazed areas were largely due to differences in land use. As is the case for most livestock exclosures, occasional trespass grazing occurred for many of the sites and wild ungulates, such as Rocky Mountain elk (*Cervus canadensis)* and mule deer (*Odocoileus hemionus*), had free access to the exclosures. Uplands were dominated by ponderosa pine (*Pinus ponderosa*) and mixed conifer forests.

Each grazed and ungrazed study reach was first delineated into pool-riffle channel units. Vegetation community composition was determined for each channel unit on both sides of the stream by calculating the percent cover of each plant species occurring in 1 x 4 m plots (N ≈ 40 plots/reach). Plots were positioned so that its center was mid-way along a a riffle or midway along a pool. Plots were then placed at the innermost edge on the green line which is the transitional point along a streambank edge where terrestrial vegetation dominates ground cover [26]. All plant species with a canopy cover 5% or more within the plot were recorded. Taxonomy largely follows that of Hitchcock and Cronquist [27].

From the plot data, we calculated species richness (number of species per experimental reach), species diversity, and similarity. Species diversity (H’) was calculated using the Shannon Index, where:

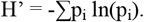

The quantity p_i_ is the proportion of cover of the ith species relative to the sum of cover for all species. We also report species diversity as the exponent of H’ which is equivalent to the number of equally common species required to produce the value of H’ [27]

The similarity between grazed and ungrazed reaches was calculated using Sorenson’s quantitative measure of similarity [27]. Similarity ranges from 0 (no species in common) to 1 (all species and their cover are identical). Similarity (C_N_) was calculated using the formula:

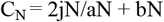

Where:

jN = sum of the lower of the two abundances (cover) of all species occurring on the grazed and exclosed reaches.

aN = The sum of plant cover in the exclosed reach

bN = The sum of plant cover in the grazed reach

The wetland species prevalence index, also referred to as the wetland score, was calculated for each grazed and ungrazed reach to determine predominance of hydrophytic (wetland) vegetation [29]. This prevalence index was computed by weighting the species cover from plots with index values for wetland indicator categories (Supporting Information Table S1). Wetland indicator values were assigned to each species using the National List of Plant Species that Occur in Wetlands ([29], Supporting Information Table S1). By assigning the composition to the wetland indictor scores we derived a wetland score for each grazed and ungrazed reach. These scores can range from 1 (all species wetland-obligate) to 5 (all species upland-obligate). The wetland prevalence index was calculated as follows:

Wetland prevalence index = ∑A_i_ W_i_ / ∑W_i_

Where:

A_i_ = abundance (cover) of species i

W_i_ = indicator index value for species i

i = species

Riparian shrub and tree composition was measured via the line intercept technique. Two transects were established along the entire length of the sampled reaches; one on each side of the creek and running along the green line. The total length of these transects in the study reaches ranged from 188 to 688m. All shrubs were identified to species. The cover of every individual shrub or tree overhanging the green line was measured regardless if there was overlap with other individuals. From these measurements, woody plant composition and streamside cover was determined.

Vegetation overstory and emergent cover over the stream was measured along each channel unit. The overstory cover can not only include streamside vegetation but also tall trees (conifers) in adjacent uplands that may function as providing stream shade. This measure utilized a concave spherical densiometer. The densiometer was taped so that there was a “V” exposing only 17 of the grid line intersections following the methods of Platts et al. [31]. Cover was measured approximately 30 cm above the surface of the water, at approximately 30 cm from each bank as well as the middle of the stream. One reading at each bank was taken and four readings were taken in the middle of the stream. The four readings in the middle included readings upstream, downstream, towards right bank and towards left bank. If emergent plants (principally Cyperaceae and Juncaceae) were present in the channel, their cover relative to the entire water surface area of the channel unit was estimated.

To statistically determine if there were differences between grazed and ungrazed treatments, the study reach was used to represent an experimental unit. Paired t-tests and the Wilcoxen sign tests were used to test for differences in prevalence indices, diversity indices and cover between grazed and ungrazed treatments [32]. Given the inherently high variation in composition, land use histories and differences in stream geomorphology, we assumed statistical differences existed when p-values were ≤ 0.10 but report the actual values. We used simple linear regression to assess the relationship between riparian vegetation variables (i.e., species diversity, individual species abundances, richness, and wetland prevalence index) with time since exclusion and channel gradient.

## Results

Riparian zones are environments of high species richness and diversity. There were 128 plant species encountered with a cover of at least 5% in one of the microplots (Supporting Information Table S3). At all sites we found significant differences in vegetation composition and structure that we attribute to livestock exclusion. The cover of native sedges (*Carex* spp.) was significantly greater (p=0.004) in ungrazed areas (Fig. 2). Forbs (broad-leaved herbs) were more prevalent ungrazed reaches (P=0.05) and shrub cover was greater in 88% of the ungrazed reaches that were >5 years old. In contrast, bare ground was higher in 63% of the grazed reaches (Table 2).

**Table 2.** Herbaceous vegetation cover (%) and bare ground (%) of riparian vegetation life forms in 11 paired exclosed (ungrazed) and grazed reaches in northeastern Oregon. Cover of herbaceous components are the means of 40 plots per experimental reach. Emergent and overstory cover is that which occurs over the stream channel.

| Cover (%) |  |  |  |  |  |  |  |  |  |  |  |  |  |
| --- | --- | --- | --- | --- | --- | --- | --- | --- | --- | --- | --- | --- | --- |
|  | p- value | Bear |  | Camas |  | Chesnimnus |  | Devils |  | Lower Swamp |  | Upper Swamp |  |
|  |  | Exclosed | Grazed | Exclosed | Grazed | Exclosed | Grazed | Exclosed | Grazed | Exclosed | Grazed | Exclosed | Grazed |
| Sedges | 0.004 | 29.6 | 18.4 | 20.6 | 17.1 | 9.2 | 1.5 | 14.7 | 0.7 | 38.7 | 4.6 | 33.8 | 4.7 |
| Rushes | 0.375 | 2.2 | 9.5 | 8.0 | 4.2 | 0.4 | 0.3 | 0.9 | 0.4 | 14.9 | 0.4 | 13.5 | 0.9 |
| Grasses | 0.156 | 23.1 | 30.3 | 37.9 | 33.9 | 33.1 | 33.0 | 41.9 | 58.1 | 39.8 | 68.1 | 35.9 | 35.2 |
| Forbs | 0.048 | 49.0 | 68.7 | 40.5 | 41.6 | 50.5 | 36.2 | 59.5 | 46.0 | 35.2 | 20.4 | 43.1 | 31.7 |
| Bare Ground | 0.129 | 25.4 | 9.6 | 25.1 | 19.7 | 14.3 | 24.1 | 4.7 | 12.2 | 1.7 | 8.0 | 0.4 | 36.2 |
| Emergent | 0.067 | 1.4 | 0.5 | 5.3 | 6.8 | 0.6 | 1.0 | 3.1 | 0.8 | 23.5 | 2.5 | 11.8 | 2.9 |
| Overstory | 0.207 | 18.0 | 19.8 | 56.8 | 3.7 | 39.8 | 48.3 | 38.7 | 10.4 | 51.5 | 78.5 | 31.2 | 35.8 |

Table 2. (continued)
| Cover (%) |  |  |  |  |  |  |  |  |  |  |  |  |
| --- | --- | --- | --- | --- | --- | --- | --- | --- | --- | --- | --- | --- |
|  | Mid Fk John Day |  | Murderers |  | Summit |  | Tex |  | Camp |  | All sites |  |
|  | Exclosed | Grazed | Exclosed | Grazed | Exclosed | Grazed | Exclosed | Grazed | Exclosed | Grazed | Exclosed | Grazed |
| Sedges | 46.6 | 12.7 | 33.4 | 29.4 | 54.7 | 20.9 | 5.7 | 4.4 | 12.7 | 10.6 | 27.2 | 11.4 |
| Rushes | 14.2 | 1.6 | 51.8 | 68.6 | 10.2 | 6.5 | 0.1 | 3.3 | 26.7 | 18.0 | 13.0 | 10.3 |
| Grasses | 37.5 | 66.5 | 42.0 | 30.6 | 11.1 | 11.1 | 41.3 | 35.6 | 52.5 | 61.9 | 36.0 | 42.2 |
| Forbs | 48.5 | 28.0 | 50.9 | 21.1 | 43.9 | 50.4 | 61.6 | 40.4 | 74.3 | 67.6 | 50.6 | 41.1 |
| Bare Ground | 15.9 | 15.4 | 4.4 | 6.0 | 10.6 | 22.9 | 1.9 | 10.9 | 0.6 | 10.0 | 9.5 | 15.9 |
| Emergent | 13.8 | 12.8 | 13.3 | 8.2 | 3.6 | 4.4 | 3.7 | 0.6 | 5.3 | 1.4 | 7.8 | 3.8 |
| Overstory | 2.7 | 0.4 | 32.4 | 13.2 | 22.1 | 4.2 | 70.3 | 67.2 | 56.8 | 43.9 | 38.2 | 29.6 |

**Figure 2.**
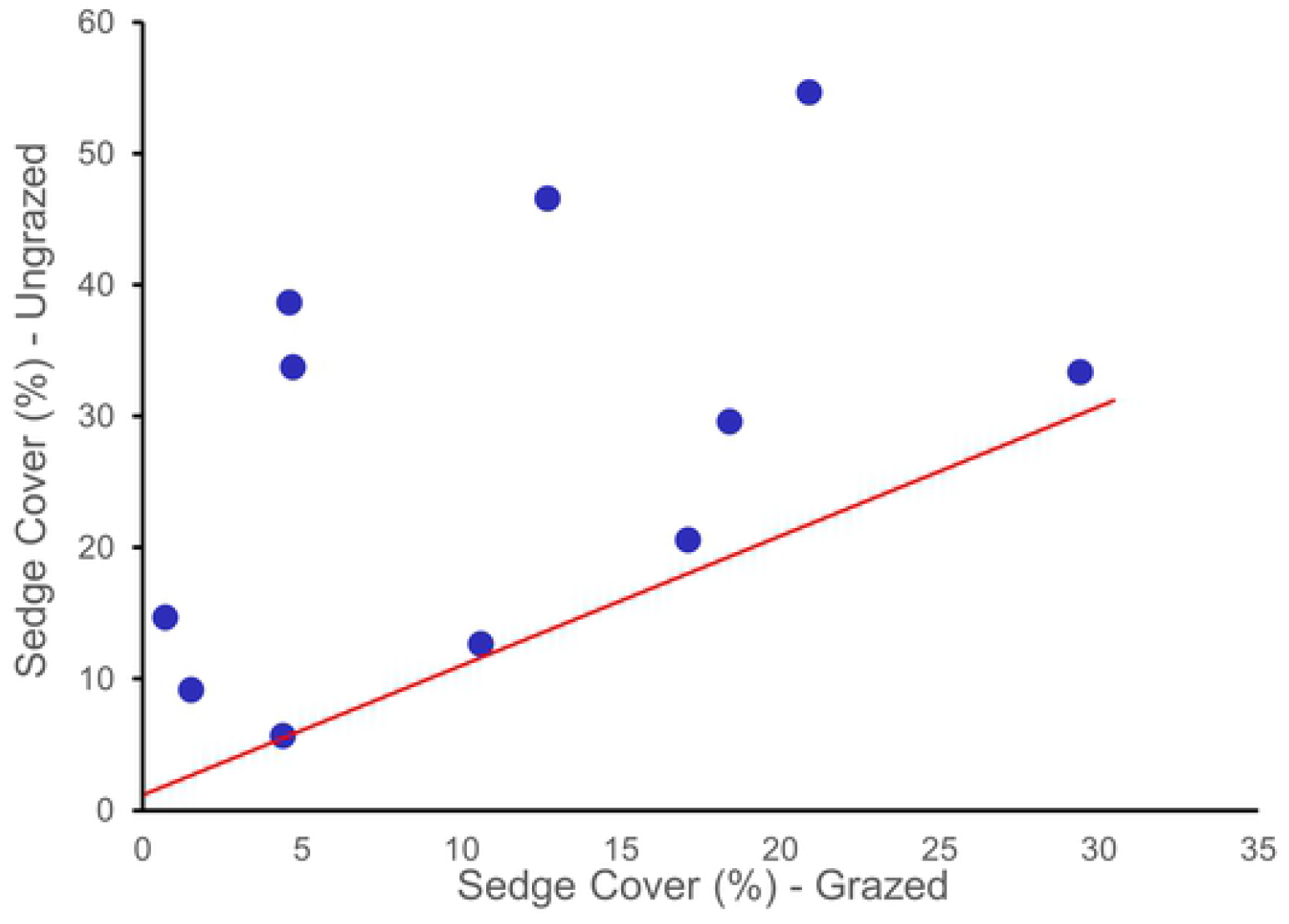
Comparison of sedge cover for 11 paired grazed and ungrazed riparian reaches in Northeast Oregon, 2000. If there was no change in the abundance of sedges between grazed and ungrazed areas the data points would be expected to fall on or near the line. Data points above the line indicate a higher abundance of sedges in ungrazed areas compared to grazed reaches. The abundance of sedges in exclosures was significantly greater P = 0.004) than in grazed areas.

Species richness (p =0 .08) and diversity(P= 0.002) was significantly greater in the ungrazed reaches (Table 3). The highest species richness was found in ungrazed reaches of Camp (S=50), Camas (S= 45), Chesnimnus (S= 45) and Devil Creeks (S=45). The greatest differences in species richness were in the heavily grazed Devil’s Creek study reach where the ungrazed portion had 17 more plant species than the grazed portion. Species diversity (exp H’) was as low 4.7 and 5.5 in the grazed reaches of the Middle Fork John Day and Murderer’s Creek, respectively. However, the ungrazed reaches of these two streams had species diversity values of 14.6 and 12.7, indicating that livestock were limiting species diversity on these sites (Table 3). Similarly, there was a large difference in the species diversity in the grazed and ungrazed reaches of Summit Creek (9.1 and 17.7, respectively). Sites with the fewest differences included the recently established Bear (2 yrs) and Camas Creek (5 yrs) sites, as well as the forested Tex Creek (23 yrs). Tex Creek was a moderately constrained reach with a dense conifer overstory canopy and light grazing regime, understory composition was most similar at Tex Creek (a similarity index of 0.73).

**Table 3.** Plant species richness (S), species diversity (H’ and exp H’), similarity (C_S_), wetland species prevalence indices (WPI), and dominant species and cover (%) for paired exclosed (ungrazed) and grazed stream reaches in northeastern Oregon.

| Site | S | H' | exp H' | C<sub>S</sub> | WPI | Dominant species (% cover) |
| --- | --- | --- | --- | --- | --- | --- |
| Bear |  |  |  |  |  |  |
| Exclosed | 40 | 3.10 | 22.10 | 0.62 | 2.82 | Poa pratensis (10), Fragaria virginiana (10), Carex pellita (9) |
| Grazed | 41 | 3.02 | 20.54 |  | 3.00 | Equisetum arvense (19), Trifolium repens (16), Poa pratensis (15) |
| Camas |  |  |  |  |  |  |
| Exclosed | 45 | 3.17 | 23.83 | 0.55 | 2.73 | Poa pratensis (19), Carex pellita (7), Juncus balticus (5) |
| Grazed | 45 | 3.11 | 22.42 |  | 2.89 | Poa pratensis (15), Trifolium repens (7), Phleum pratense (7) |
| Chesnimnus |  |  |  |  |  |  |
| Exclosed | 45 | 2.80 | 16.48 | 0.41 | 2.60 | Equisetum arvense (30), Poa pratensis (22), Salix fragilis (18) |
| Grazed | 36 | 2.61 | 13.54 |  | 3.28 | Poa pratensis (27), Pseudotsuga menziesii (11), Trifolium repens (11) |
| Devils |  |  |  |  |  |  |
| Exclosed | 45 | 2.90 | 18.09 | 0.35 | 2.68 | Myosotis scorpiodes (32), Agrostis stolonifera (15), Alnus incana (10) |
| Grazed | 28 | 2.17 | 8.76 |  | 3.57 | Poa pratensis (37), Trifolium repens (19), Phleum pratense (17) |
| Lower Swamp |  |  |  |  |  |  |
| Exclosed | 36 | 2.74 | 15.50 | 0.34 | 2.50 | Agrostis stolonifera (25), Carex utriculata (20), Alnus incana (18) |
| Grazed | 34 | 2.02 | 7.50 |  | 3.04 | Alnus incana (52), Poa pratensis (49), Agrostis stolonifera (10) |
| Upper Swamp |  |  |  |  |  |  |
| Exclosed | 35 | 2.92 | 18.46 | 0.42 | 2.46 | Agrostis stolonifera (18), Erigeron philadelphicus (15), Alnus incana (13) |
| Grazed | 30 | 2.67 | 14.37 |  | 2.53 | Juncus balticus (13), Poa pratensis (9), Trifolium repens (9) |
| Mid Fk J. Day |  |  |  |  |  |  |
| Exclosed | 32 | 2.68 | 14.62 | 0.34 | 2.11 | Carex pellita (30), Solidago lepida(15), Deschampsia cespitosa (15) |
| Grazed | 23 | 1.55 | 4.71 |  | 3.71 | Poa pratensis (64), Solidago lepida (11), Carex pellita (10) |
| Murderers |  |  |  |  |  |  |
| Exclosed | 32 | 2.54 | 12.70 | 0.59 | 1.86 | Juncus balticus (44), Poa pratensis (23), Carex utriculata (21) |
| Grazed | 21 | 1.74 | 5.45 |  | 1.63 | Juncus balticus (67), Carex nebrascensis (27), Poa pratensis (24) |
| Summit |  |  |  |  |  |  |
| Exclosed | 37 | 2.88 | 17.74 | 0.38 | 1.95 | Carex nebrascensis (19), Carex pellita (16), Carex utriculata (14) |
| Grazed | 29 | 2.21 | 9.12 |  | 3.32 | Trifolium longipes (31), Poa pratensis (10), Carex pellita (9) |
| Tex |  |  |  |  |  |  |
| Exclosed | 36 | 2.63 | 13.86 | 0.73 | 2.76 | Alnus incana (54), Poa pratensis (16), Symphyotrichumfoliaceus (11) |
| Grazed | 39 | 2.53 | 12.59 |  | 2.92 | Alnus incana (40), Poa pratensis (14), Equisetum arvense (8) |
| Camp |  |  |  |  |  |  |
| Exclosed | 51 | 3.01 | 20.21 | 0.63 | 2.86 | Poa pratensis (34), Alnus incana (30), Juncus balticus (24) |
| Grazed | 41 | 2.80 | 16.50 |  | 3.15 | Poa pratensis (51), Trifolium longipes (26), Juncus balticus (16) |
| P-value | 0.08 | 0.002 | 0.004 |  | 0.01 |  |

The changes in species richness and diversity reflect significant changes in species composition between the grazed and ungrazed reaches. Plant species composition were least similar (0.34-0.38) between reaches of Devils, Lower Swamp, Middle Fork John Day, and Summit Creeks. These were all comparatively low gradient reaches where sedges and relatively hydric species were found in greater abundance in ungrazed areas. The relatively young age of some of these exclosures demonstrates that species composition shifts of the herbaceous component can occur in a relatively short period of time following cattle exclusion. We found the most similar composition between grazed and ungrazed sites to be in the forested reaches with a relatively high gradient (i.e., Tex and Camp Creek) and at some of the recently established exclosures. Combining all sites in the analysis we found no strong relationships (r^2^ =0.01) between gradient and shifts in sedge composition. There were also no strong relationships between channel or valley gradients with the shifts in the parameters of diversity.

Wetland-obligate and facultative wetland species were found in greater abundances in exclosures. Wetland species prevalence indices were consistently and significantly lower (p = 0.01) in the ungrazed reaches compared to the grazed reaches (Fig. 3, Table 3). This indicates increased hydric conditions and soil moisture available for riparian plant use following cessation of grazing. Combining all sites, we found a statistically significant increase in native sedge (*Carex* spp.) abundance (largely wetland-obligate and facultative-wetland species (Fig. 2) with concomitant declines in the abundance of the facultative exotic grass Kentucky bluegrass (*Poa pratensis*)(Fig 4). The only site where we did not find a shift towards wetlands species prevalence was at Tex Creek (a partially constrained forested reach; Fig.3).

**Figure 3.**
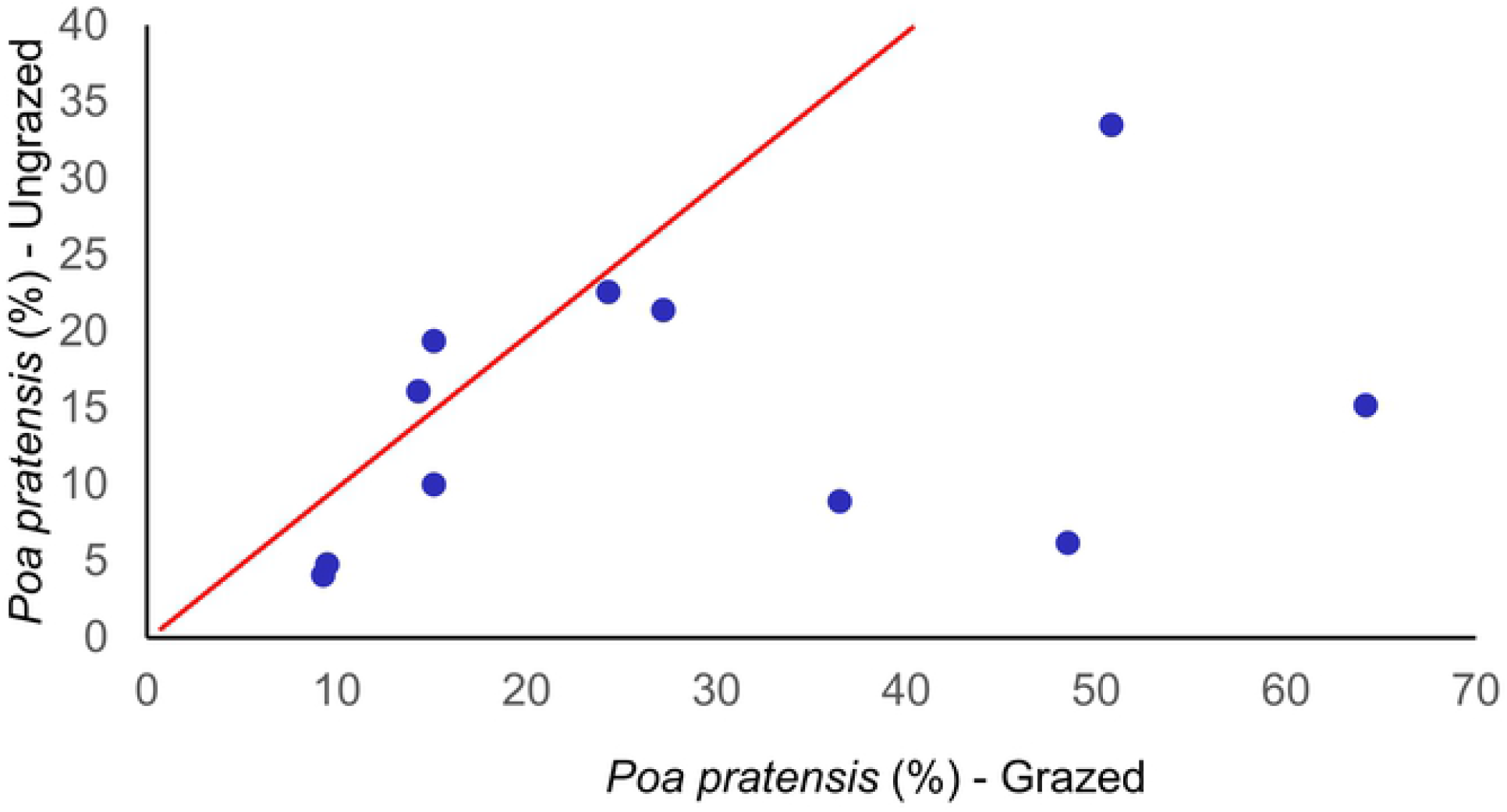
The relationship of wetland prevalence index for 11 paired grazed and ungrazed riparian reaches in Northeast Oregon. If there was no difference in the index between grazed and fenced areas the data points would be expected to fall on or near the line. Data points below the line indicates a greater abundance of wetland species in ungrazed areas compared to grazed reaches. There was a significance difference (P =0.01) in the wetland indicator scores of grazed and ungrazed areas.

**Figure 4.**
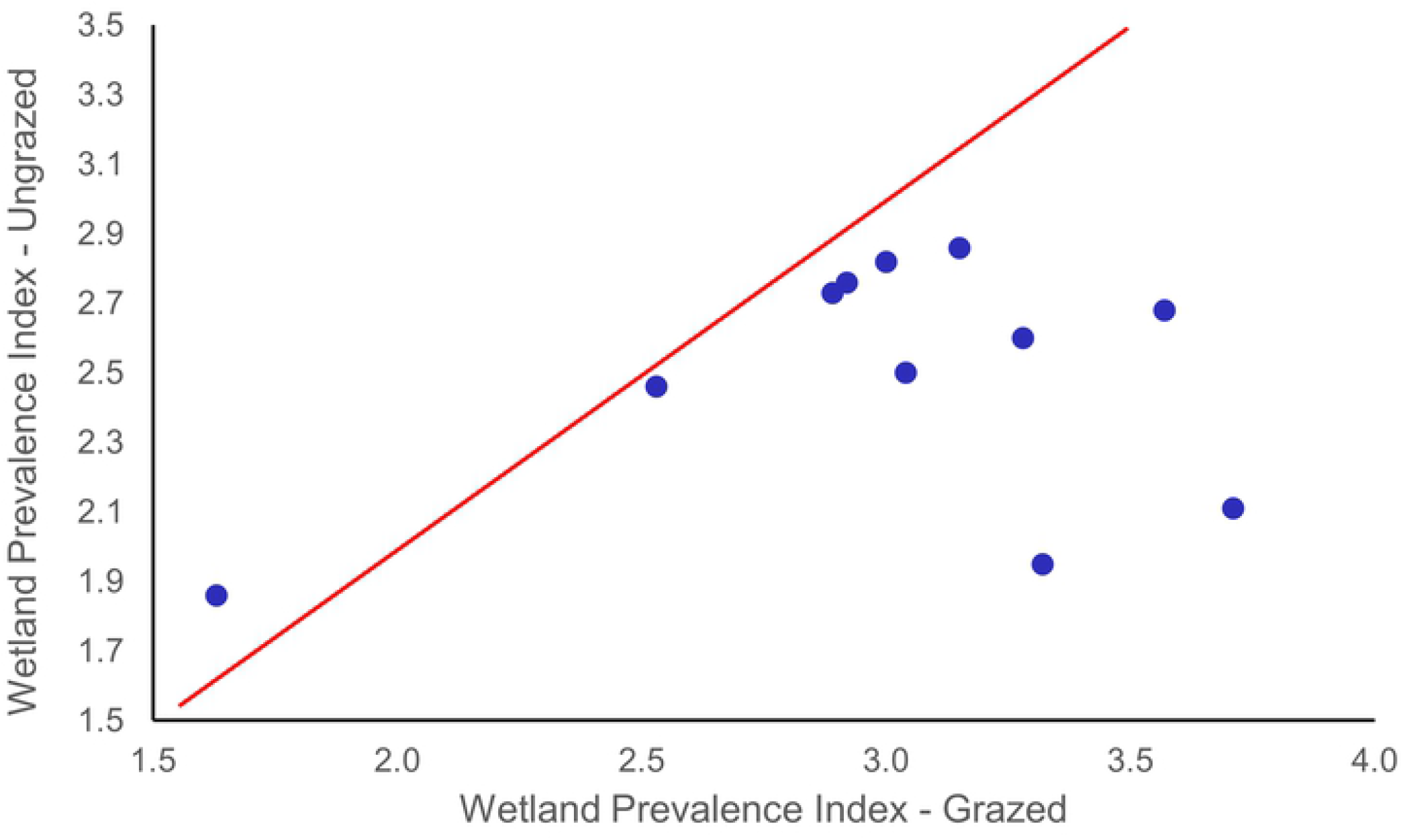
The relationship of the cover of an exotic grass, Kentucky bluegrass (P*oa pratensis*) adapted to herbivory, within 11 paired grazed and ungrazed riparian reaches in Northeast Oregon 2000. If there was no difference in the abundance of this species between grazed and exclosed areas the data points would be expected to fall on or near the line. Data points below the line indicates a lower abundance of *P. pratensis* in ungrazed reaches compared to grazed reaches. The abundance of Kentucky bluegrass in was significantly less (p = 0.03) in ungrazed areas.

The greatest shifts in wetlands species abundances were in low-gradient reaches (e.g., Middle Fk John Day, Summit Creek, and Lower Swamp Creek). For example, at Summit Creek, wetland scores were 41% lower in ungrazed reaches (1.95) compared to grazed reaches (3.32); the streambank in the ungrazed reach was dominated by wetland-obligate and facultative wetland species while streambank in the grazed reach was dominated by facultative and facultative upland species. For example, *Carex* species (wetland obligates) were most common in ungrazed reaches (55% cover) while facultative species *Poa pratensis* and *Trifoliun longipes* dominated the grazed reaches (50% cover; Table 3). Hydric species such as *Carex* spp., *Glyceria* spp., and *Scirpus microcarpus* were more abundant in the ungrazed reaches while species more adapted to grazing and drier conditions (e.g., *Poa pratensis, Phleum pratense, Trifolium* spp., *and Taraxacum officinale*) were usually more abundant in grazed reaches (Fig. 4). There was a mean 49% decrease in the cover of the exotic grass *Poa pratensis* in ungrazed reaches (p = 0.03, Fig 4).

Stream reaches also varied in their potential to support tree or shrub-dominated communities (Table 6). Streamside cover of woody vegetation ranged from 0.7% in the low gradient, meadow-dominated Middle Fk John Day to 129% in the relatively steep-gradient, forested Tex Creek. The most abundant woody species was Thin-leaf alder (*Alnus incana)*, which was present in 10 of the 11 sites. A total of 6 willow species (*Salix* spp.) were encountered in the study but never in great abundance. Overall, we encountered 28 shrub and tree species in the study with *Salix* spp. being present at all sites.

Total shrub cover was higher in ungrazed reaches for six of the 11 study sites; riparian-obligate shrub cover was greater in ungrazed reaches for seven of the 11 study sites. Cover differences can be largely explained by the age of the exclosure. Of the four study streams where cover was equal or less in ungrazed compared to grazed areas, 3 of them were less than 5 years old. In 7 of 8 study streams where exclosures were >5 years in age, shrub cover was greater in the ungrazed areas. The greatest differences in shrub cover were in the oldest 4 exclosures (>20 years). For example, woody vegetation cover at Summit creek (22 years) was 26 and 6% in ungrazed and grazed reaches, respectively, while woody vegetation cover at Camp creek (36 years) was 74 and 35%, respectively (Table 4). This finding indicates that 20 years or more of livestock exclusion may be required to allow significant recovery of woody vegetation. The benefits, ecosystem services, and values of the stream reaches excluded from livestock increase through time and may not be fully realized until decades after exclusion.

**Table 4.**
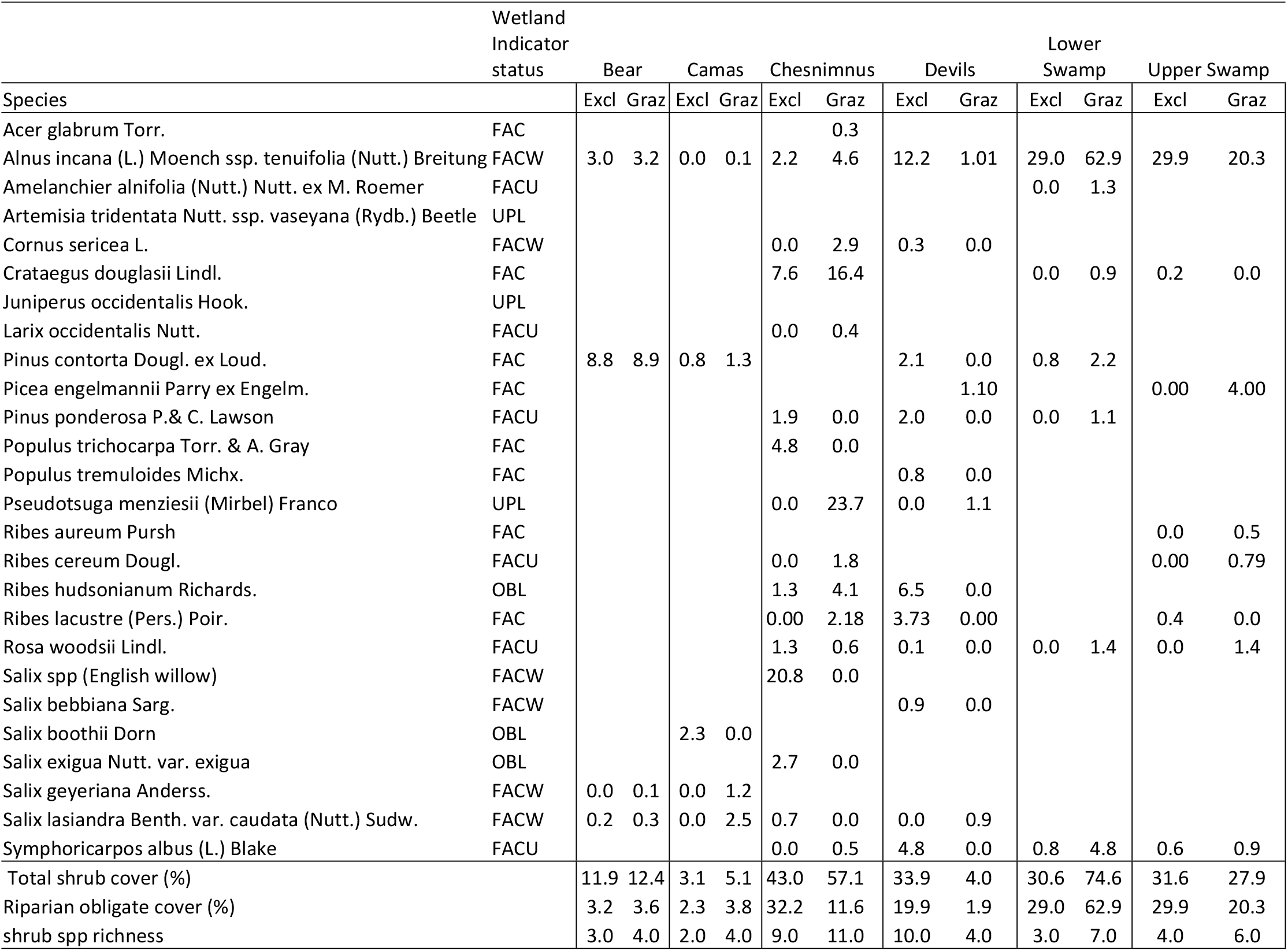
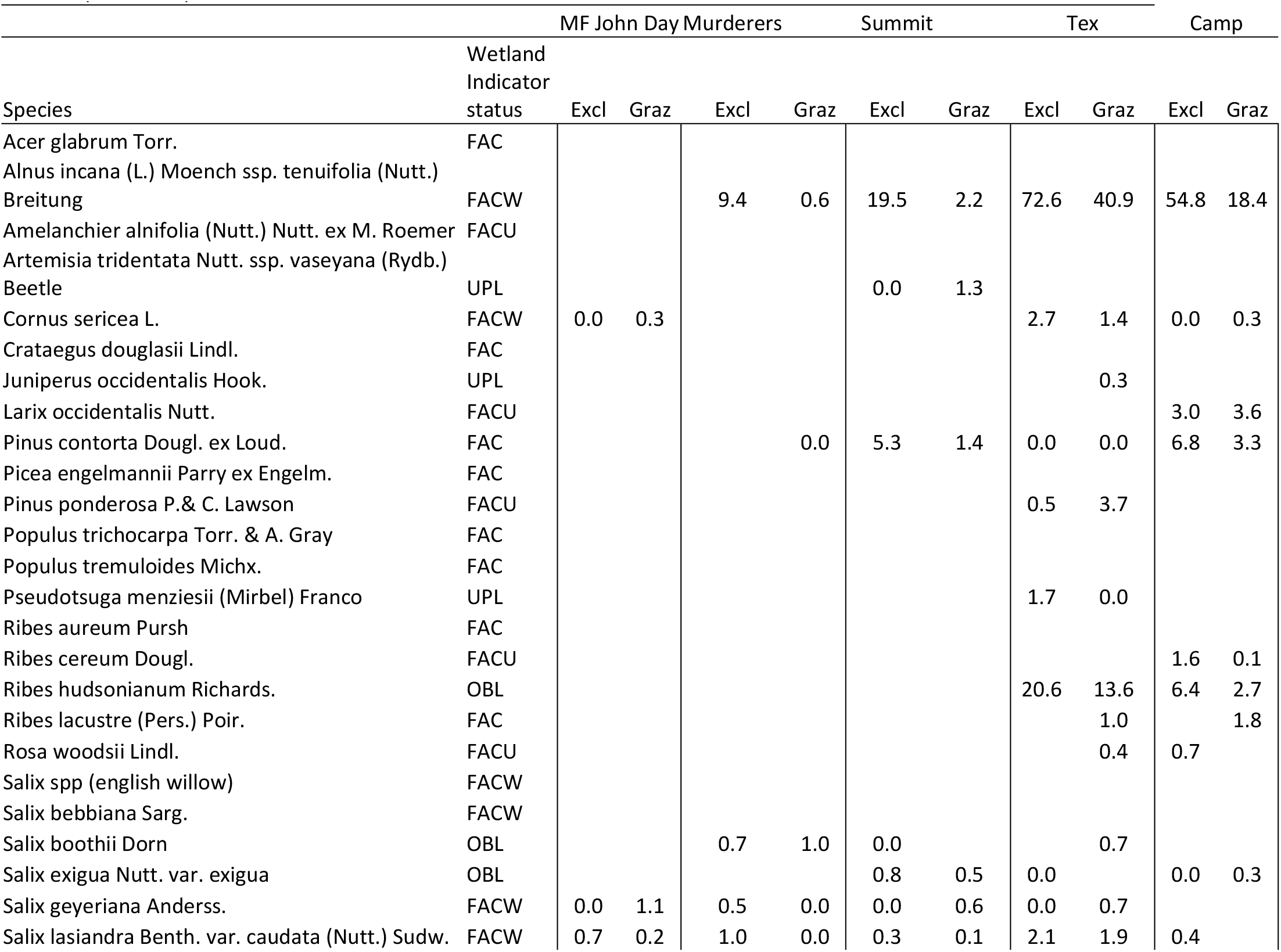
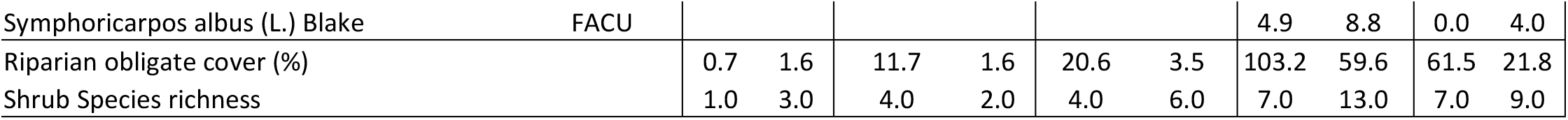
Shrub and tree cover (%) along the streambank (greenline) of selected stream study reaches in northeastern Oregon.

While differences in the structure and abundance of shrub species was often related to age since exclusion, we did not find strong relationships between time since exclusion and either changes in species richness or species diversity. This suggests that potential increases in species richness, diversity, and wetland prevalence indices were highly variable between sites, perhaps due to differences in geomorphology, edaphic conditions, and land use history.

## Discussion

Ecological restoration of riparian habitats is defined as the reestablishment of predisturbance riparian functions and related chemical, biological, and hydrological characteristics [33]. Passive restoration is defined as the halting of those activities that are causing degradation or preventing recovery [24] Reviews of many instream restoration and enhancement projects throughout the western U.S. clearly reveal that passive restoration, especially the cessation of livestock grazing was the critical first step in successful riparian restoration programs [17, 18, 20, 34]. The apparent resilience of riparian vegetation composition reflected in the significant increases in wetland obligates and species diversity coupled with declines in exotic species and decreases in bare ground exemplifies the positive outcome of this passive restoration approach.

Cattle grazing appears to have maintained the presence of some exotic dominants at the exclusion of native species in most study reaches. In the absence of grazing, native graminoids and dicots increased in abundance while there was a concomitant decline in the exotic Kentucky bluegrass and Dutch Clover (*Trifolium repens*) was a dominant herbaceous species in 6 of the sampled stream reaches but was much reduced in adjacent exclosures (Table 3). Kentucky bluegrass is a grazing-tolerant nonnative species that has invaded rangelands in the United States [35] and is currently a dominant in many riparian zones across the western USA. Kentucky bluegrass may reduce the genetic diversity of other species through habitat fragmentation as well as ecosystem species diversity and, therefore, resilience during future environmental stress events [35]. The prevalence of Kentucky bluegrass may partially explain the lower species richness and diversity in the grazed riparian reaches. We found that there was a decline in Kentucky bluegrass dominance in the ungrazed reaches (Fig. 3) with a concomitant increase in species richness and diversity (Table 3). Schulz and Leininger [36] also found a decrease in Kentucky bluegrass in ungrazed riparian zones compared to grazed sites.

Willows, thin leaf alder, and black cottonwood (*Populus trichocarpa*) are important features of western riparian ecosystems and have multiple functional roles that influence biological diversity, water quality/quantity, and aquatic/terrestrial food webs and habitat. While shrub response was most apparent in this study with long term absence of grazing (>10 years), the rapid inherent resilience of these species has been quantified. For example, after two years in the absence of livestock on Meadow creek, Northeastern Oregon, significant increases in height, crown area, crown volume, stem diameter, and biomass were measured for willows, black cottonwood, and thin-leaf alder [21]. In addition, Brookshire et al. [22] reported that even relatively light levels of domestic livestock grazing, when coupled with wild ungulate browsing in riparian zones diminished both plant structure and reproduction of riparian willows.

Regardless of the differences in the manner and intensity in which livestock were grazed in the 11 sampled stream reaches, we found increases in wetland species and decreases in exotic species. This is similar to conclusions of a review by Elmore and Kauffman [12] that reported livestock exclusion was the most effective approach to restoring riparian ecosystems..

We hypothesized wetland-obligate and facultative-wetland species would increase in abundance in sites protected from grazing (exclosures). We found a significant 24% shift in the wetlands species prevalence index (WPI) towards wetland species in ungrazed sites (Table 3). The trends in compositional change between grazed and ungrazed reaches were similar regardless the individual grazing approaches at each site. Comparing intact and degraded stream reaches in northeast Oregon, Toledo and Kauffman [37] reported a 29% shift in WPI (1.82 to 2.45) with a loss in wetland obligate species and a concomitant shift to species more adapted to drier environments in degraded sites. Coles-Ritchie et al. [20] also reported a statistically significant shift in wetlands species abundance in exclosed compared to grazed riparian reaches in western riparian zones. Differences in the WPI would suggest that livestock activities have altered the vegetation as well as those environmental and edaphic conditions that determine which species occupy a site.

The shifts in composition due to exclusion may also be reflective of physical changes in soil properties associated with cessation of grazing. Kauffman et al. [19] found that soil bulk density was significantly lower, and soil pore space and soil organic matter was higher in exclosed riparian meadows. The mean infiltration rate for exclosed dry meadows was 13-fold greater than in grazed dry meadows (142 vs. 11 cm/h, respectively), and for wet meadows the mean infiltration rate in exclosures was 2.3 times greater than in grazed sites (80 vs. 24 cm/h, respectively). Livestock removal was found to result in significant changes in soil, hydrological, and vegetation properties that, at landscape scales, would likely have great effects on stream channel morphology, water quality, and the aquatic biota. For example, Kauffman et al. [19] estimated that under saturated conditions, a hectare of wet meadows with the pore space measured in the exclosed wet-meadow communities would contain 121,000 L/ha (121 tonnes/ha) more water in only the surface 10 cm of soil than those in the grazed wet-meadow communities.

The composition of vegetation in the ungrazed riparian areas suggest a number of synergistic effects occur when livestock are excluded. In addition, to vegetation composition shifts, Kauffman et al. [19] reported that total belowground biomass (TBGB), consisting of roots and rhizomes) in dry meadows dominated by Kentucky Bluegrass was over 50% greater in exclosures (1105 and 652 g/m^2^ in the exclosed and grazed sites, respectively). In exclosed wet meadows dominated by sedges (*Carex*-spp), the TBGB was 62% greater in ungrazed compared to grazed sites (2857 and 1761 g/m2, respectively). However, the Kauffman et al. [19] study may be underestimating true shifts in below ground biomass associated with livestock exclusion. As we found shifts in composition from facultative species (Kentucky bluegrass) to wetland-obligate species (sedges.), relevant comparisons may be that of the root mass in grazed dry meadows to that of exclosed wet meadows (i.e., a potential increase in root mass of 156 % associated with exclusion).

The source of much of the energy, carbon and nutrients of headwater streams originates from streamside vegetation [38] and both riparian meadows and forested reaches represent an important source of allochthonous materials for aquatic systems [39]. During peak flows when streamside communities are inundated, plant materials from meadow-dominated reaches become a source of organic C and nutrients to the stream. Ungrazed sites likely have high levels of organic inputs into aquatic systems for at least four reasons: (1) there is no removal of streamside vegetation via herbivory thus aboveground biomass is relatively high; (2) there is an increase in species of higher productivity (e.g., sedges. compared to Kentucky bluegrass); (3) there are dramatic increases in root mass; and (4) there is an increase in shrub and tree canopy and the concomitant increase in litterfall. Increases in species richness and diversity also suggest there is likely an increase in the timing and composition of allochthonous inputs (associated with the increase in plant species diversity).

Shifts in ecosystem structure associated with livestock exclusion are not necessarily limited to vegetation or surface soils. Vegetation response to the exclusion of livestock by fencing (e.g., increased vegetation cover and structure, as well as reduced bare ground) are important factors associated with channel morphology. For example, Magilligan and McDowell [5] reported that livestock exclusion resulted in significant stream channel adjustments such that channels in ungrazed reaches were narrower, deeper, and had more pool area than the channels in grazed reaches. The vegetation composition within ungrazed exclosures, as measured in this study, suggests an increased connectivity of riparian vegetation with their associated aquatic system.

While livestock exclusion was associated with a significant increase in sedges, broadleaved herbs, species richness, diversity and the wetlands species prevalence, there was not a strong relationship with time since exclusion. These data suggest that some sites respond more quickly than others, which is consistent with the concept of variable recovery trajectories after livestock exclusion [25].

Differences in species composition between grazed and ungrazed riparian reaches may reflect the cumulative effects of cattle grazing species that are less adapted to herbivory (e.g., willows, cottonwoods, and other keystone vegetation species) coupled with soil trampling that lowers infiltration rates and water-holding capacities of riparian soils. Channel degradation associated with bank trampling [5] and the concomitant loss of root mass associated with herbivory [19, 37] further results in drier conditions, and thus shifts in vegetation composition toward species adapted to drier environments. This suggests that cumulative livestock impacts exacerbate the increases in temperatures and changing hydrological dynamics associated with climate change that are resulting in drier conditions [40]. Dwire et al. [41] also reported that the functionality of many riparian areas has been compromised by water diversions and livestock grazing, which reduces their resilience to additional stresses that a warmer climate may bring. Removal of grazing began to reverse these effects, suggesting that cessation of livestock utilization of riparian zones is a viable approach to climate change adaptation for these important ecosystems.

## Acknowledgments

Funding for this study was provided by the Bonneville Power Administration to Oregon State University and the University of Oregon. Additional funding was provided by the Ecosystem Restoration Research Fund of the Oregon State University Foundation. We wish to thank personnel of the Oregon Department of Fish and Wildlife, the Malheur, Umatilla, and Wallowa-Whitman National Forests and all the private landowners who participated in this study. Without their generous assistance in allowing us to conduct research on their lands, as well as their inputs in approaches to land management, and the history of the study sites, this study would not have been possible. We particularly are indebted to them for their foresight into the need for restoration of fish, wildlife, and their habitats. We also thank Dr Barb Wilson for assistance with taxonomic nomenclature.

